## Supplementary material for "Diversified regulation of circadian clock gene expression following whole genome duplication": S1 appendix

### OG1v0000277 CKId/e

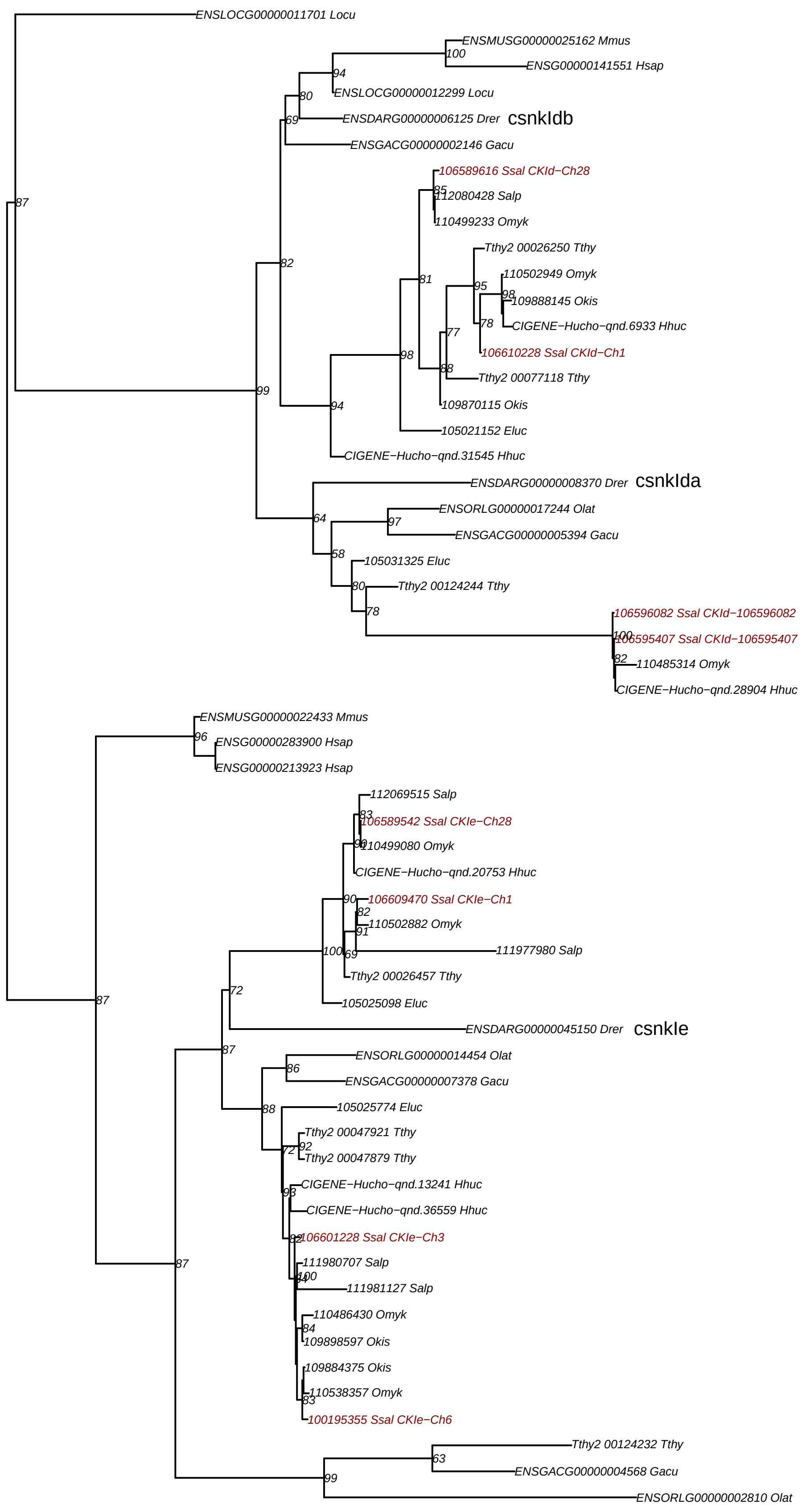


Mmus - Mouse
Hsap - Human
Locu - Spotted gar
Drer - Zebrafish
Gacu - Stickleback
Olat - Medaka
Eluc - Northern pike
Tthy - Grayling
Hhuc - Danube salmon
Salp - Arctic charr
Okis - Coho salmon
Omky - Rainbow trout
Ssal - Atalntic salmon

### **OG1v0000304** CRY

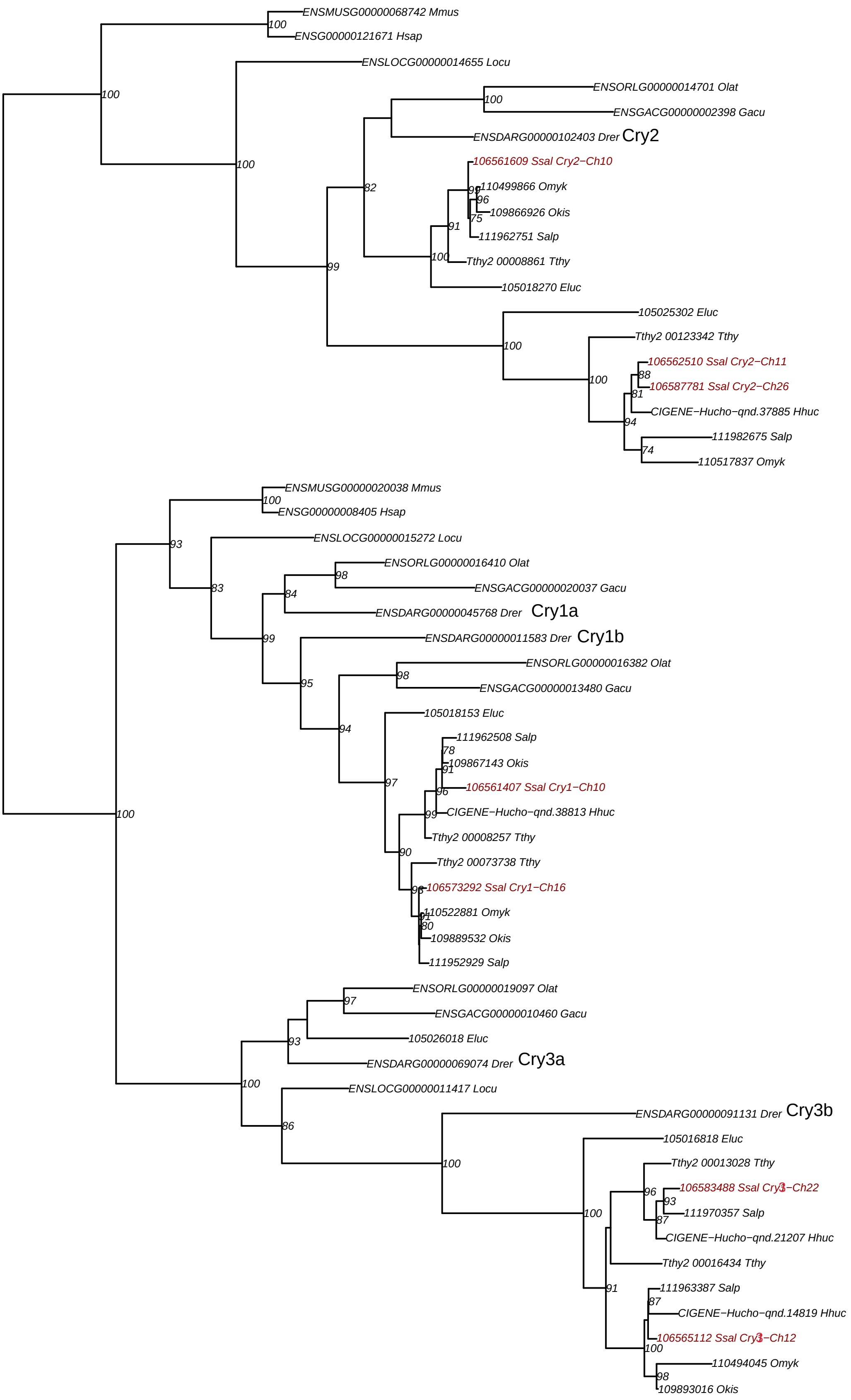

**OG1v0000681** CLOCK/NPAS

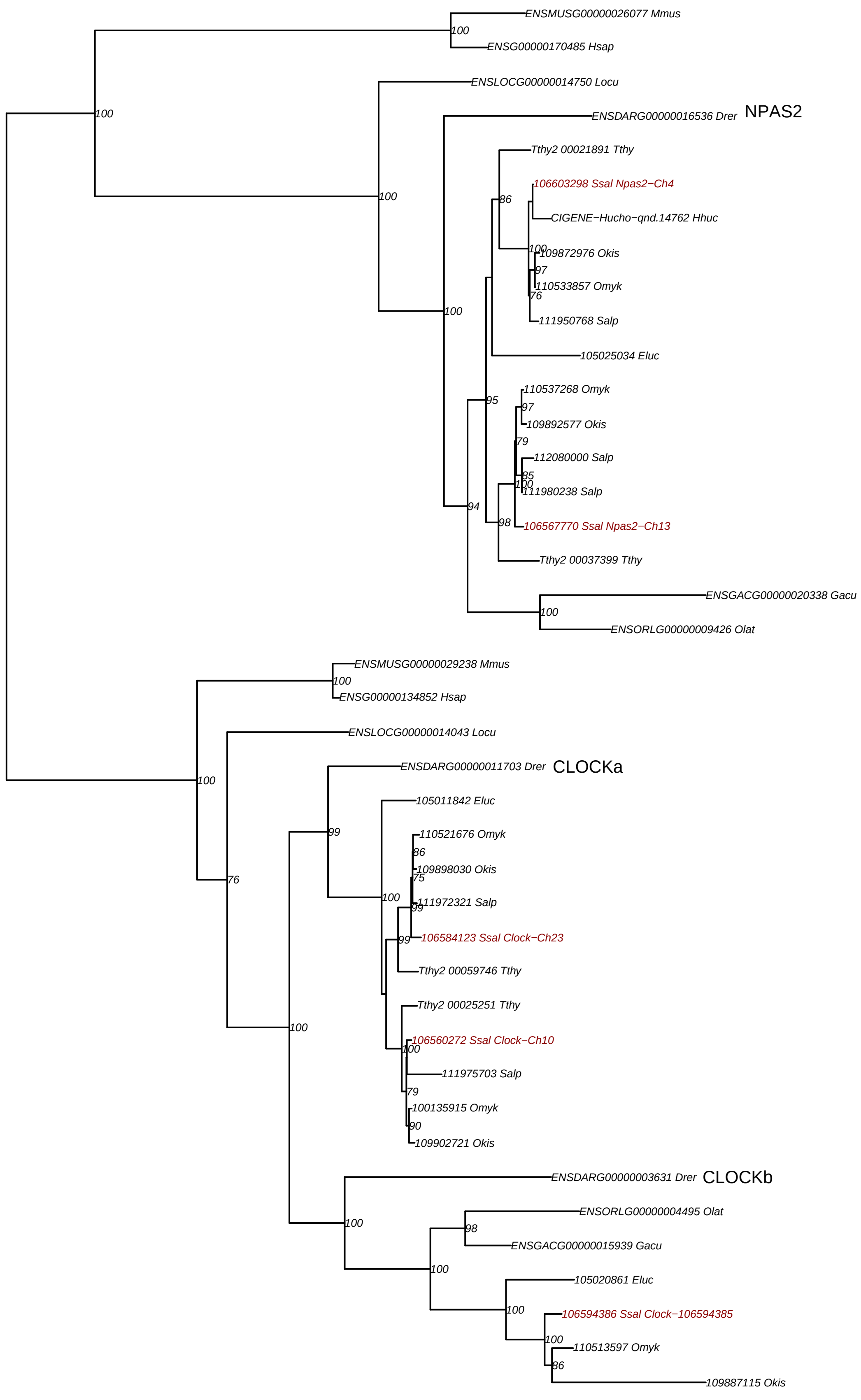

### OG1v0001103 TEF

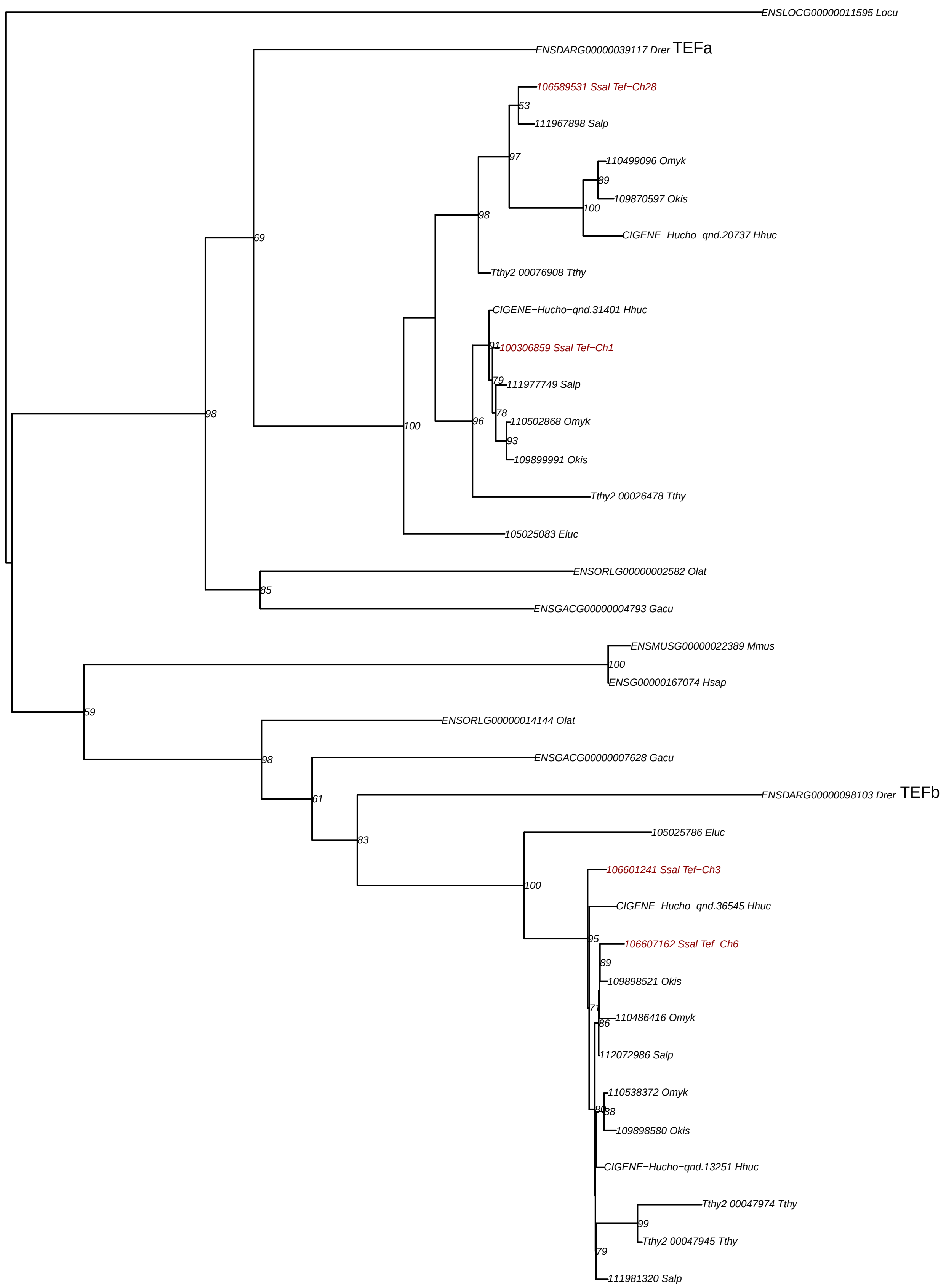

### **OG1v0001835** DBP

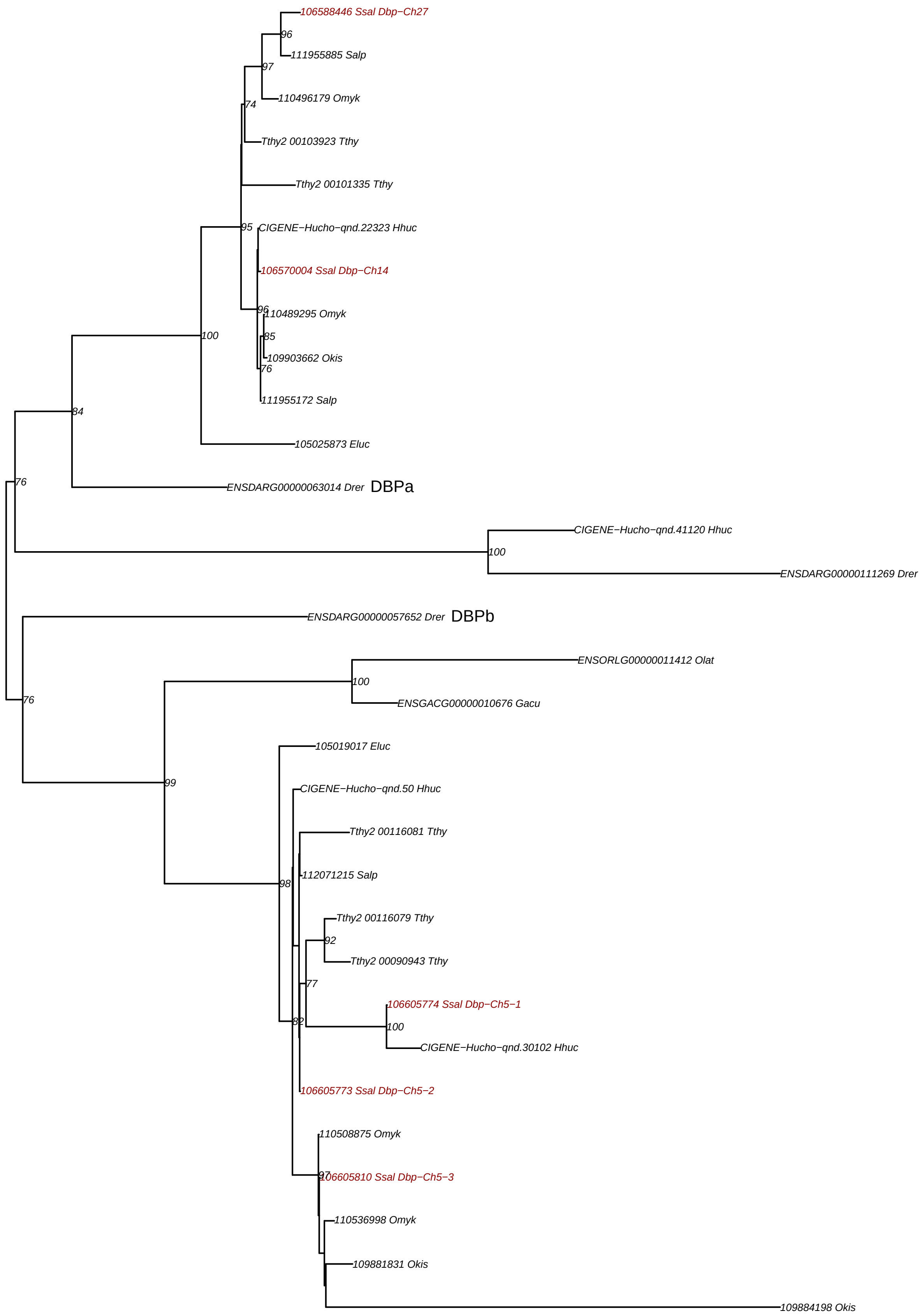

### OG1v0002483 PER1

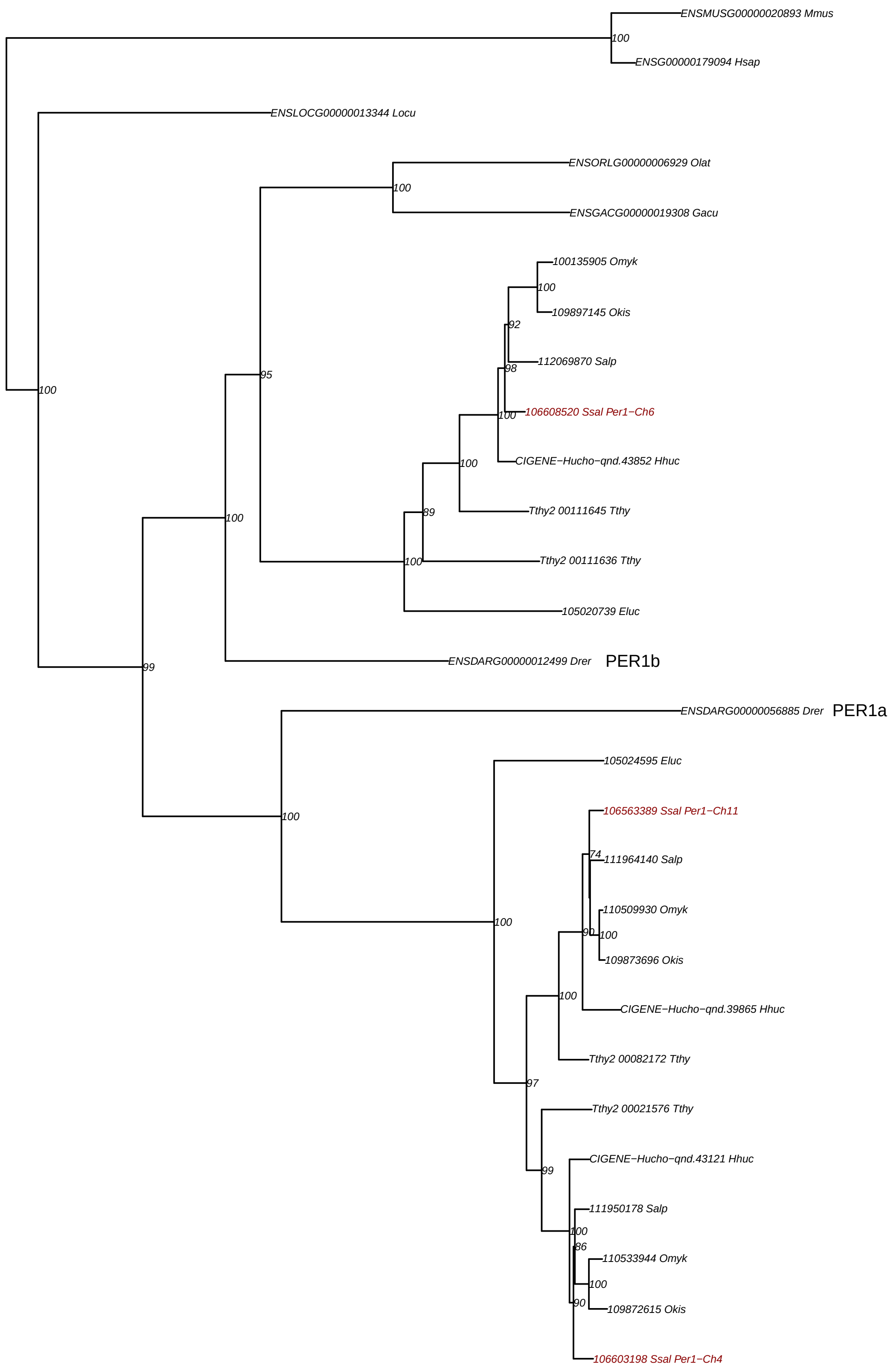

### **OG1v0002849** ARNTL1

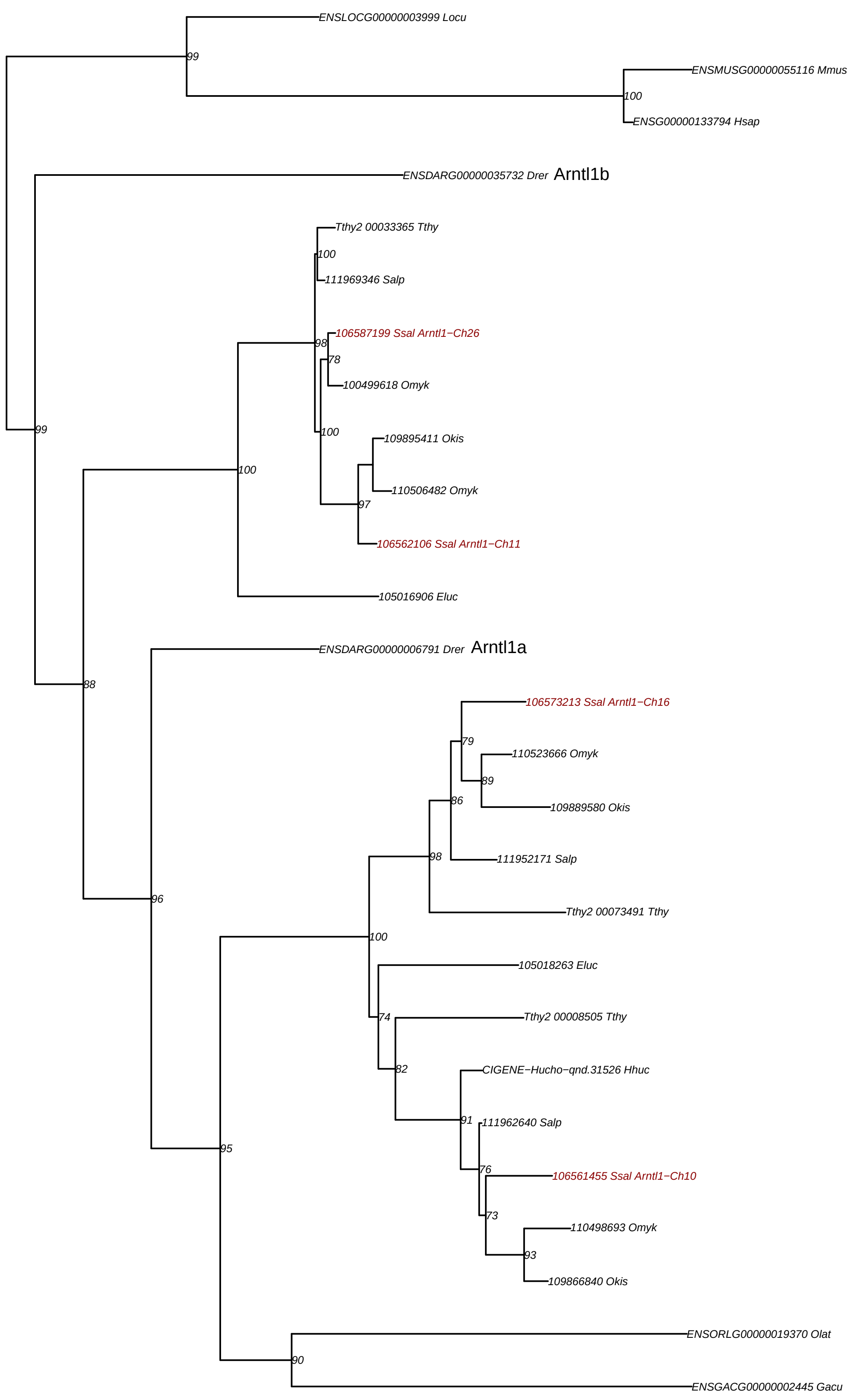

### OG1v0003243 HLF

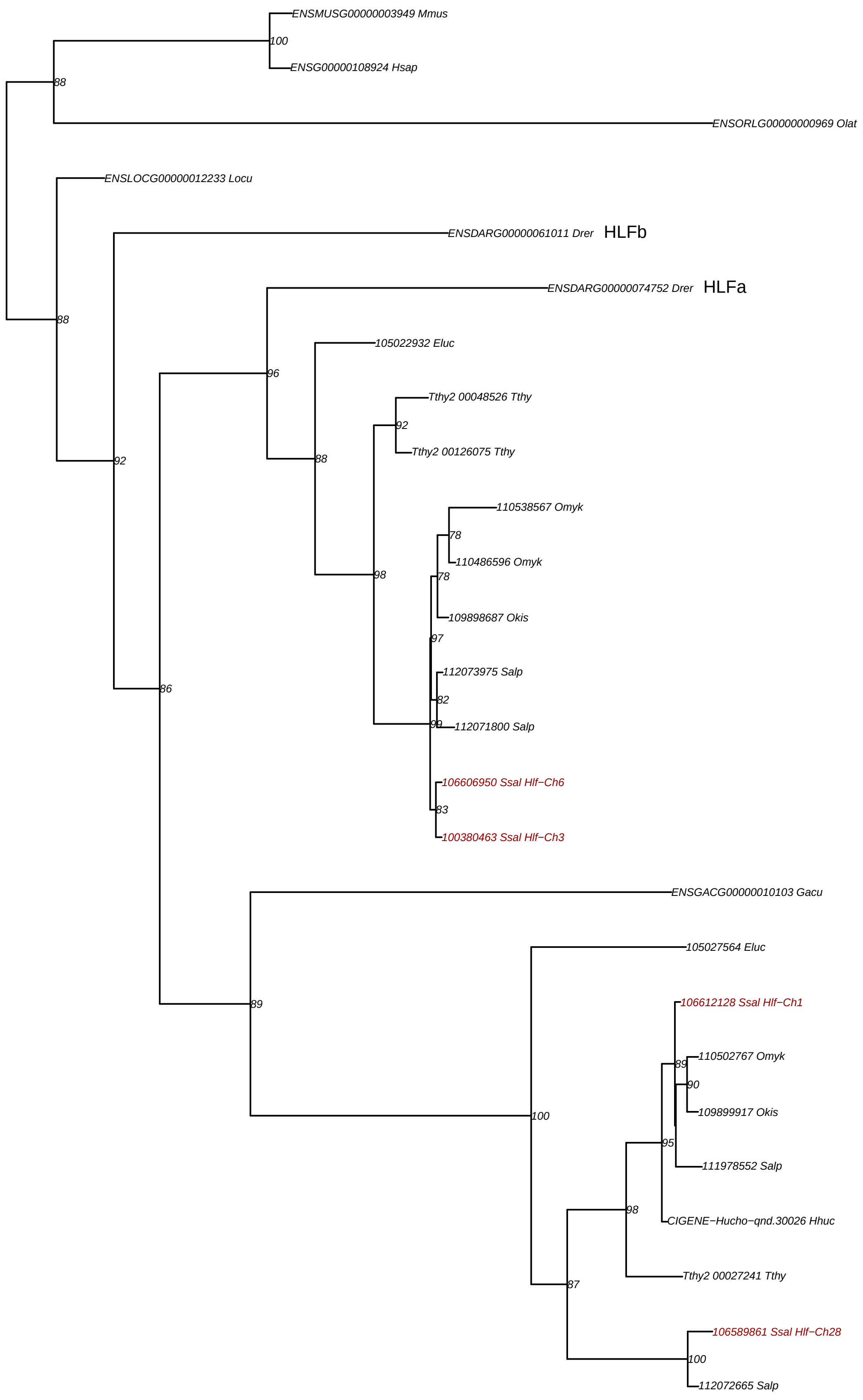

### **OG1v0003721** RORA

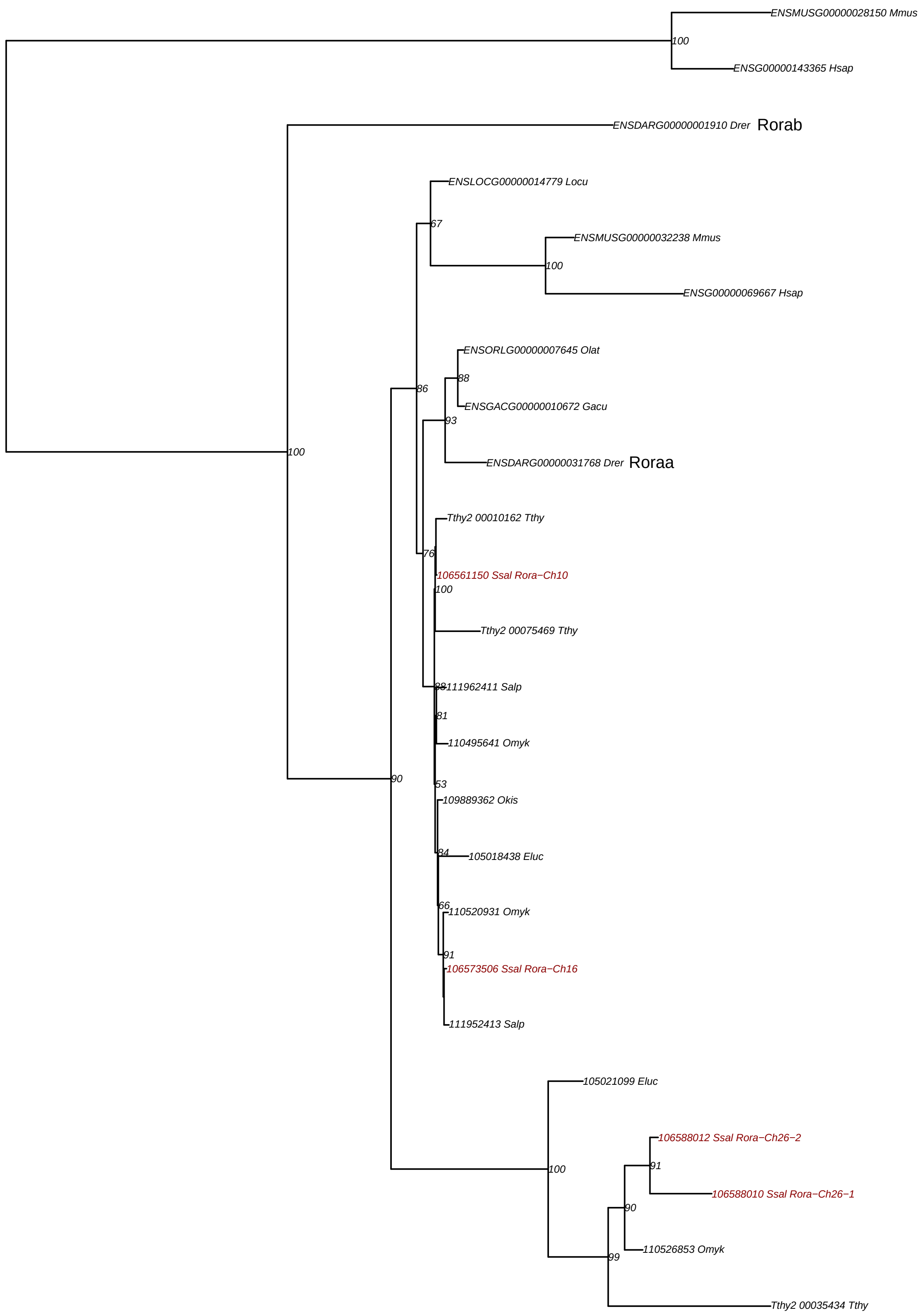

### OG1v0003904 PER2

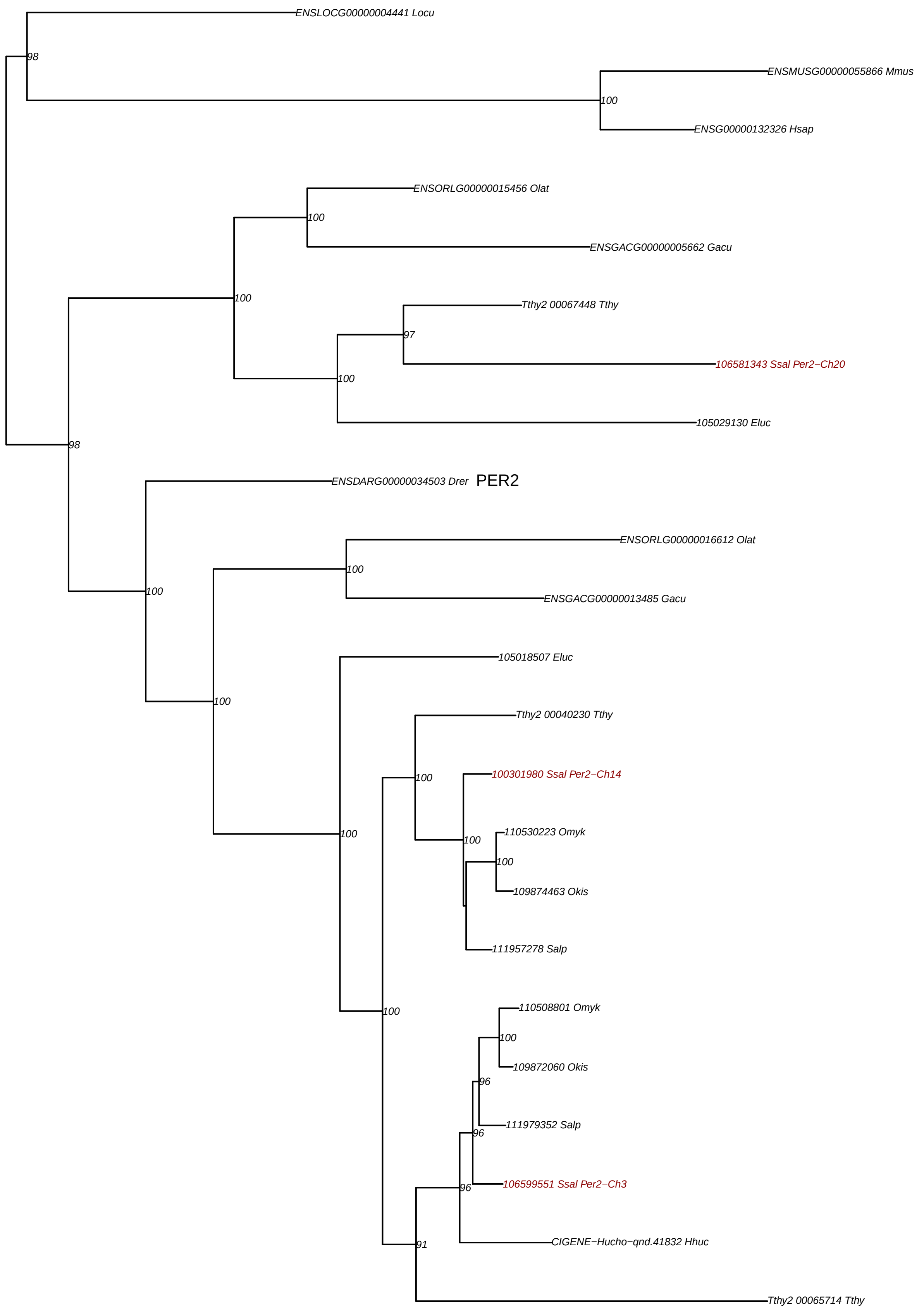

# **OG1v0007203** NR1D2b

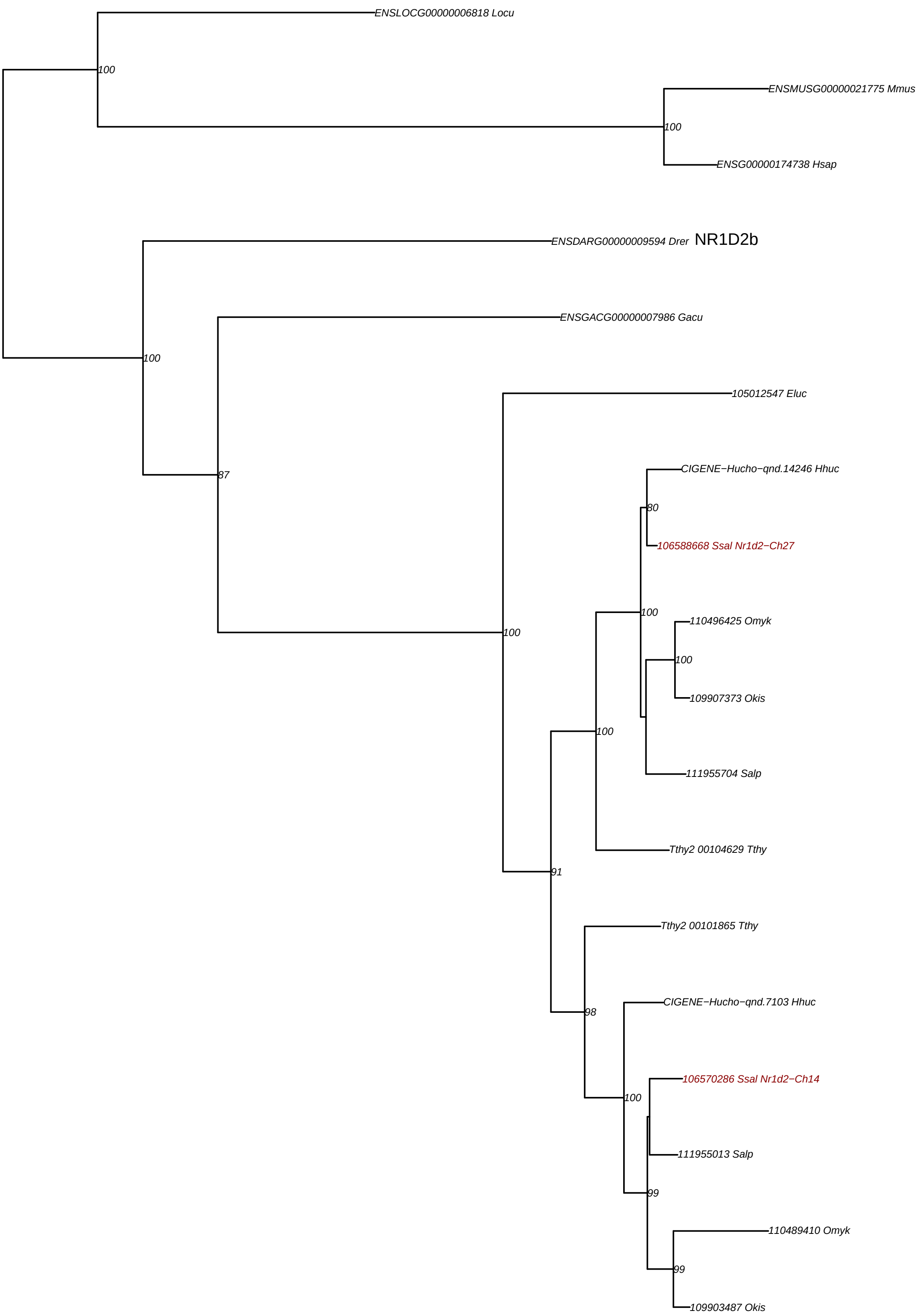

### **OG1v0007372** RORB

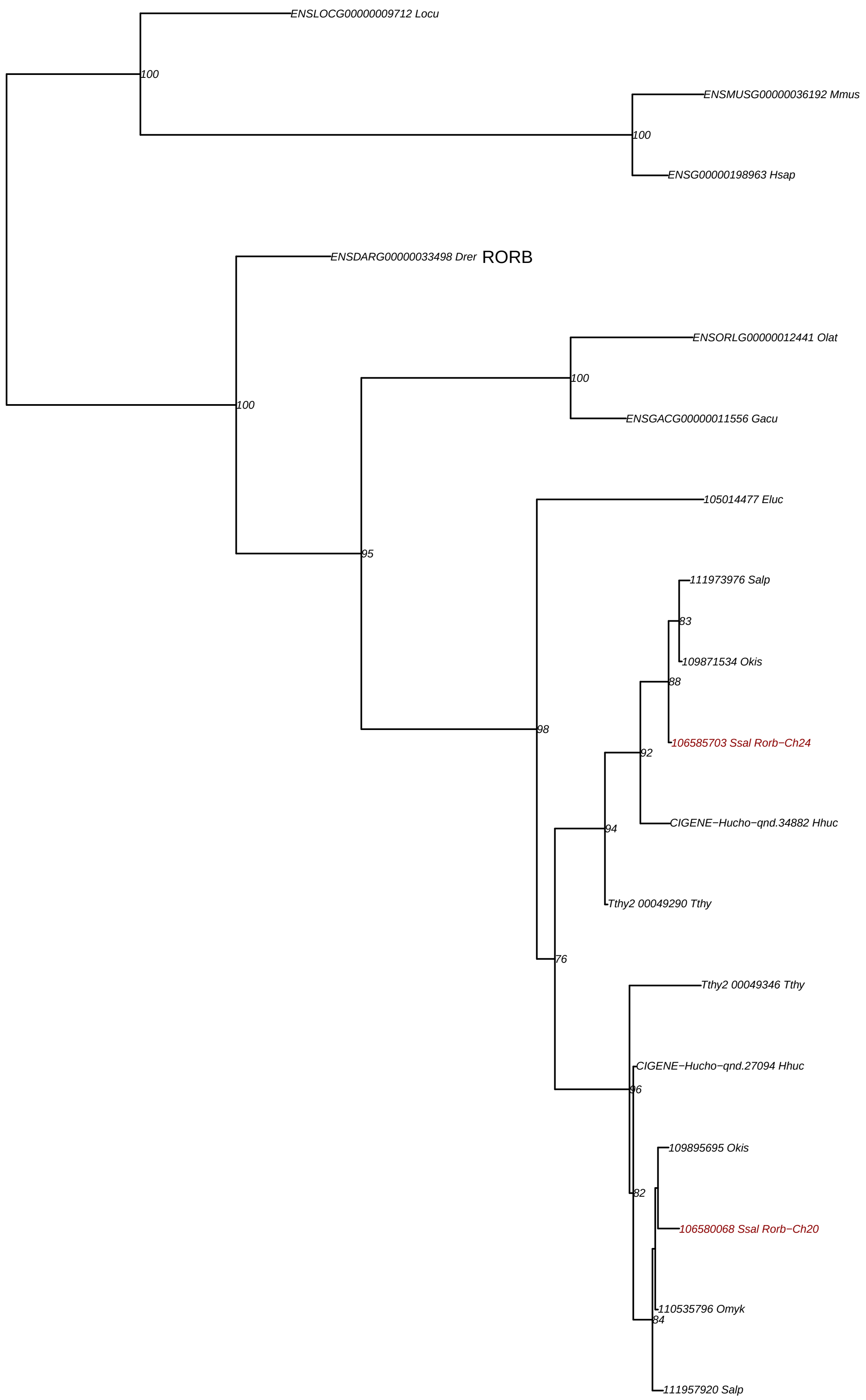

### **OG1v0007602** CIART

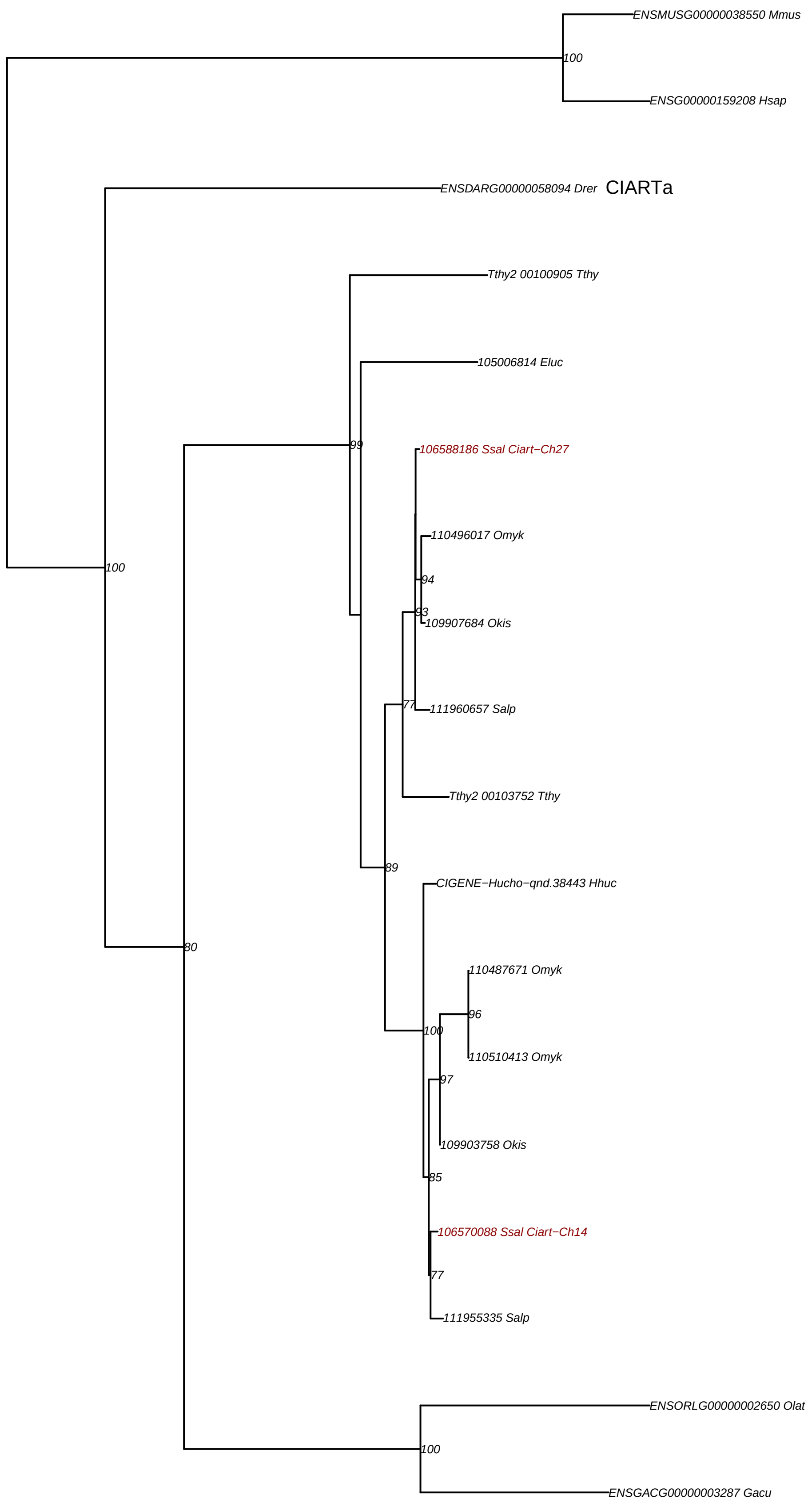

# **OG1v0010710** NR1D2a

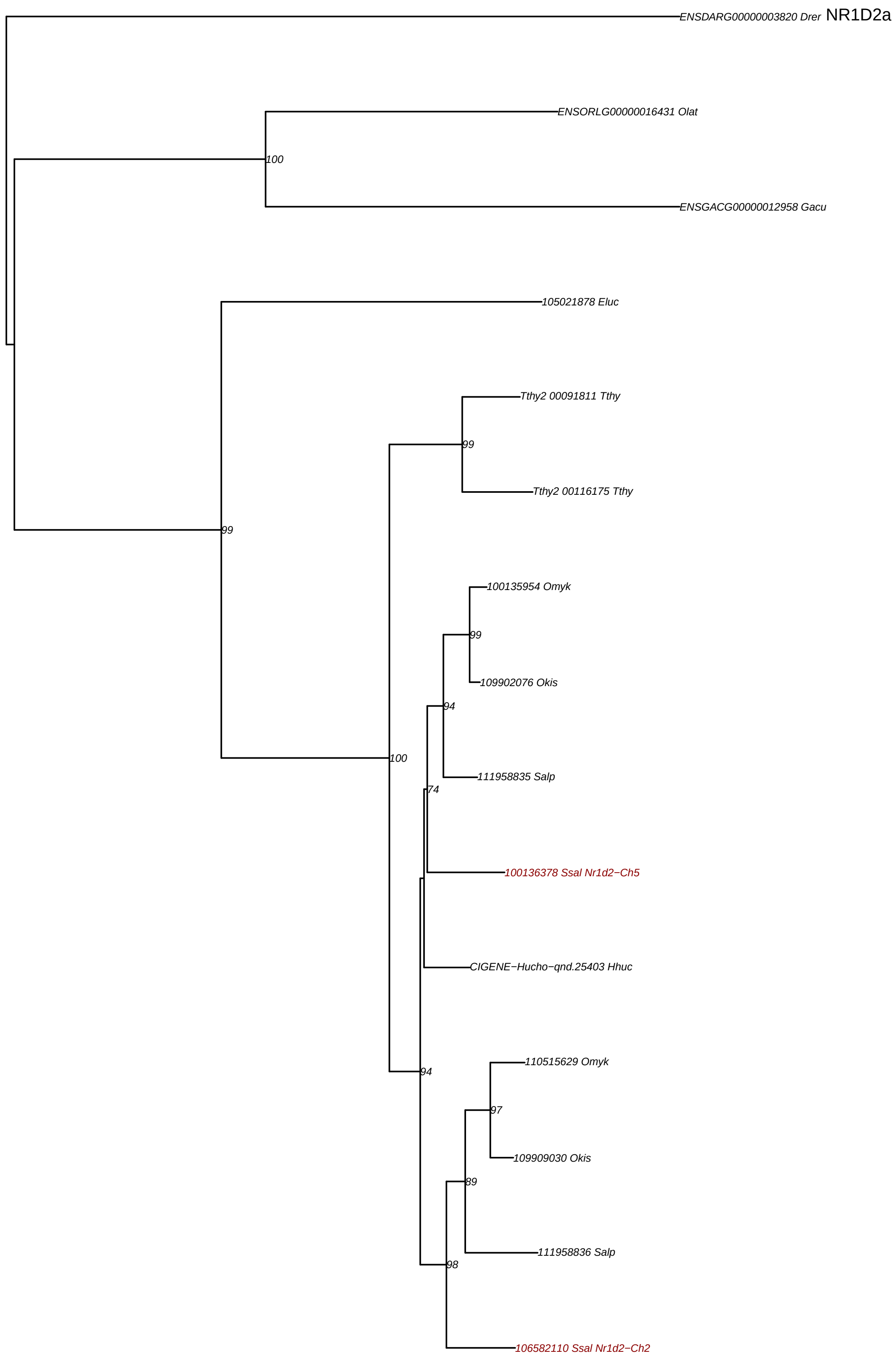

# **OG1v0011513** NR1D1

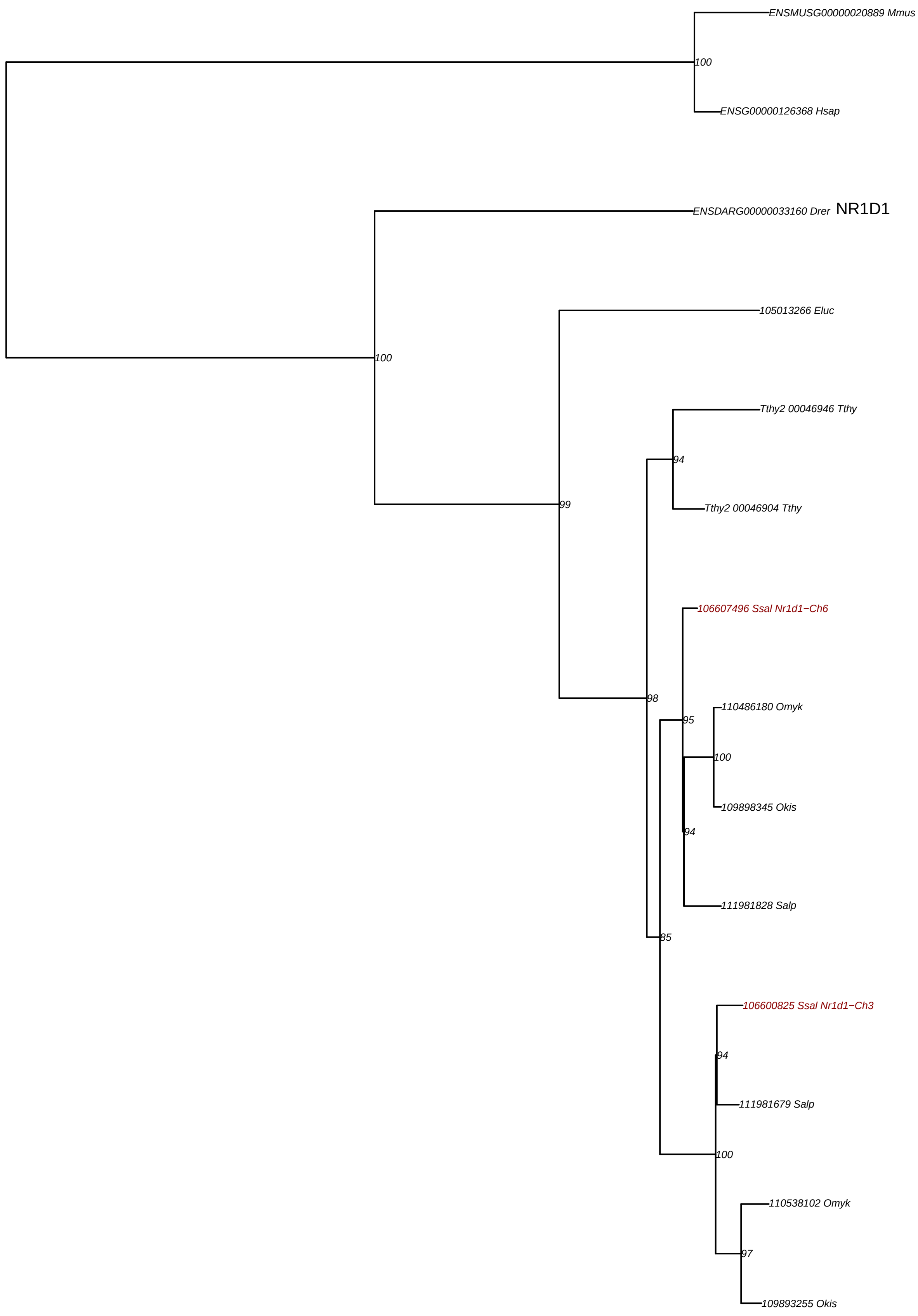

### **OG1v0013005** RORC

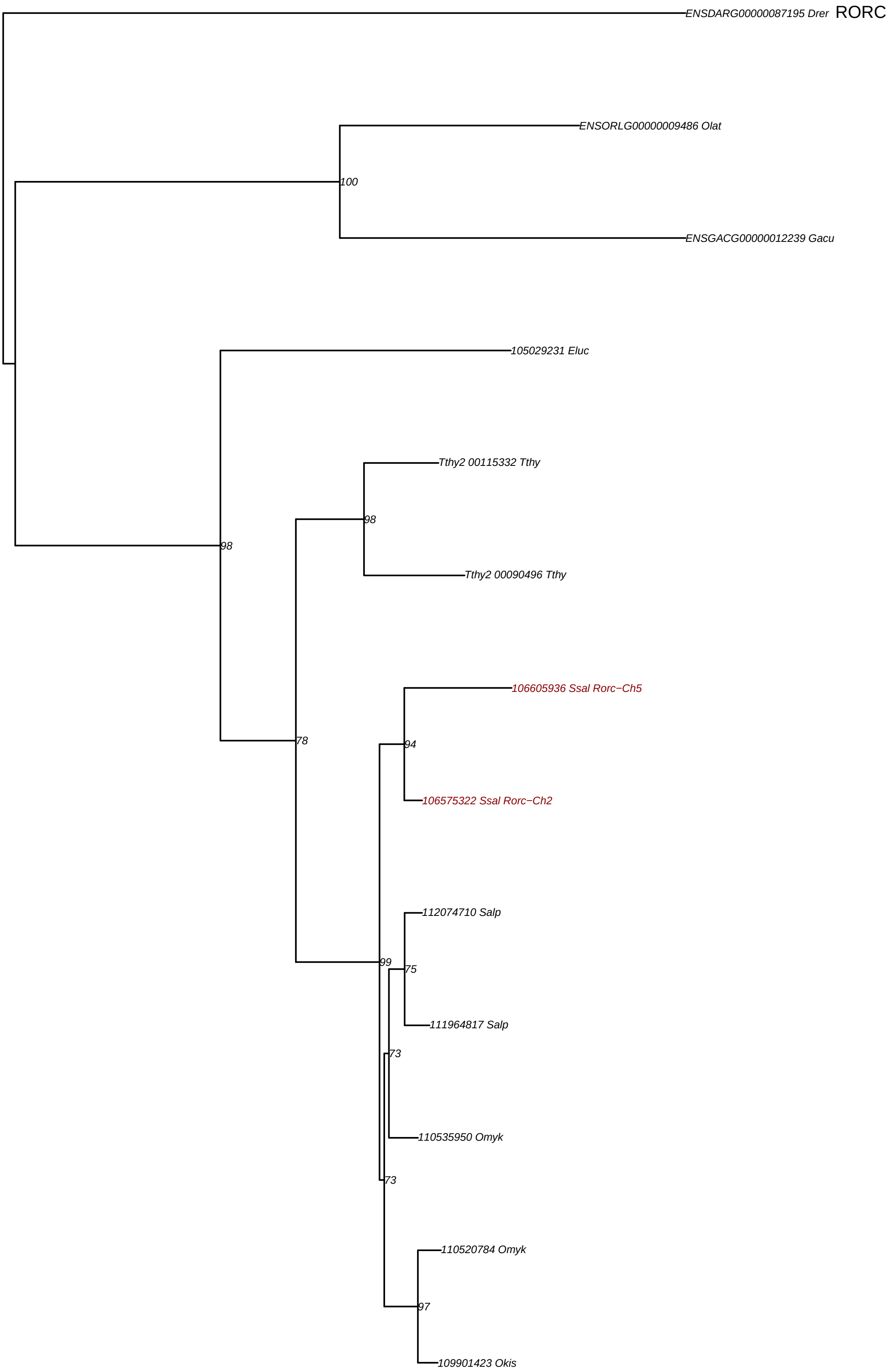

**OG1v0015055** CRY

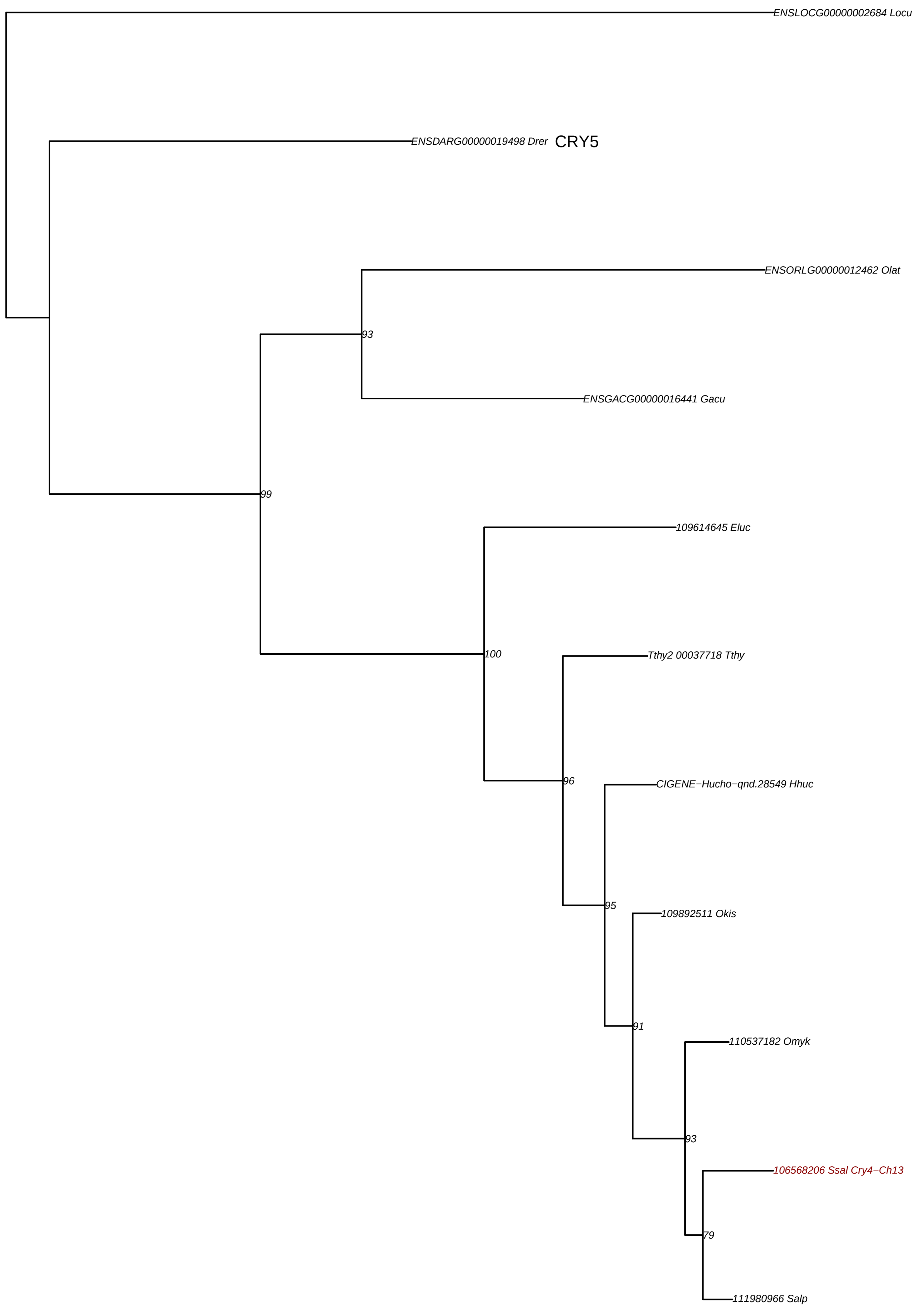

### OG1v0016141 PER3

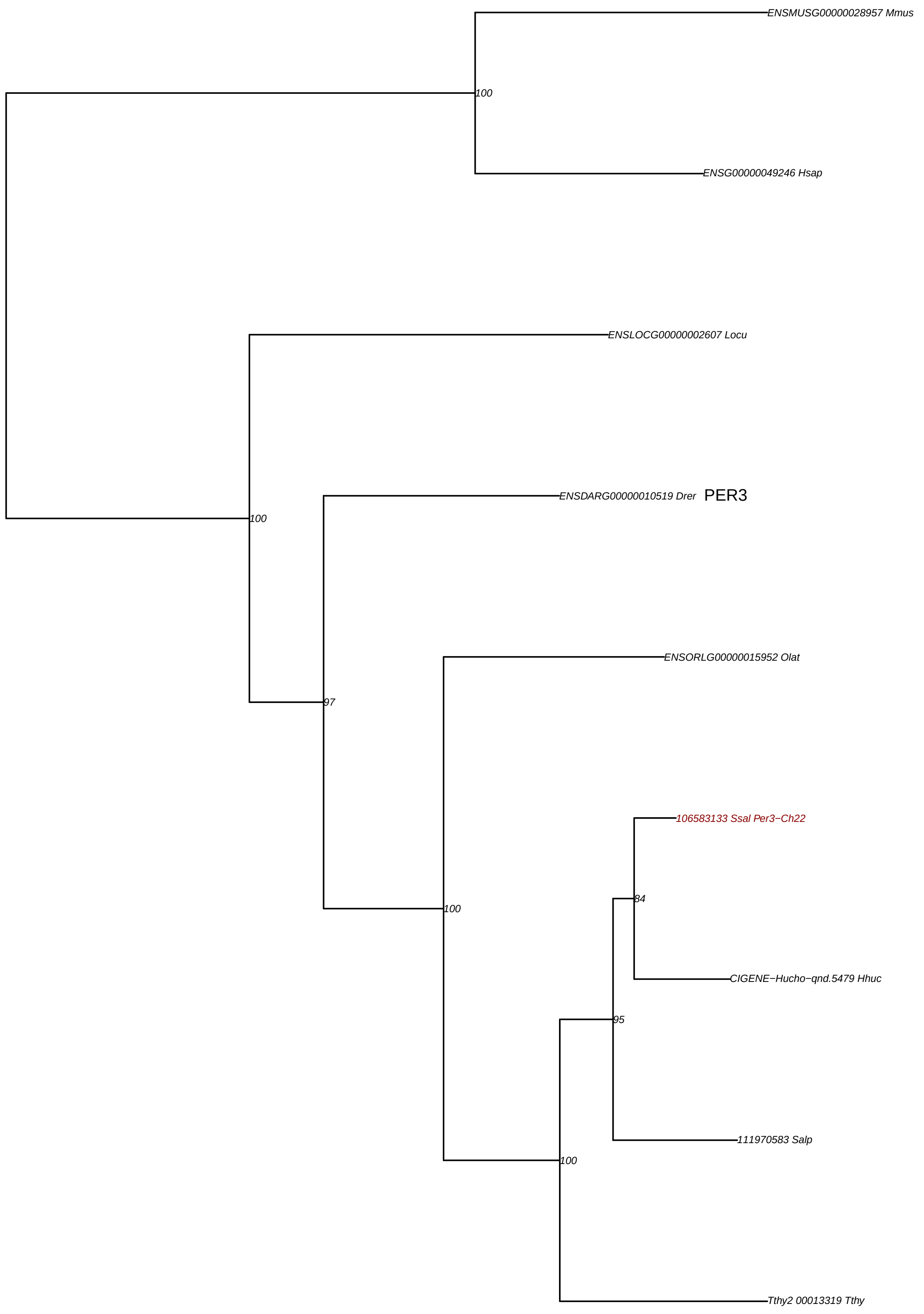
