## Supplementary figures and images for "Diversified regulation of circadian clock gene expression following whole genome duplication"

### S2 appendix

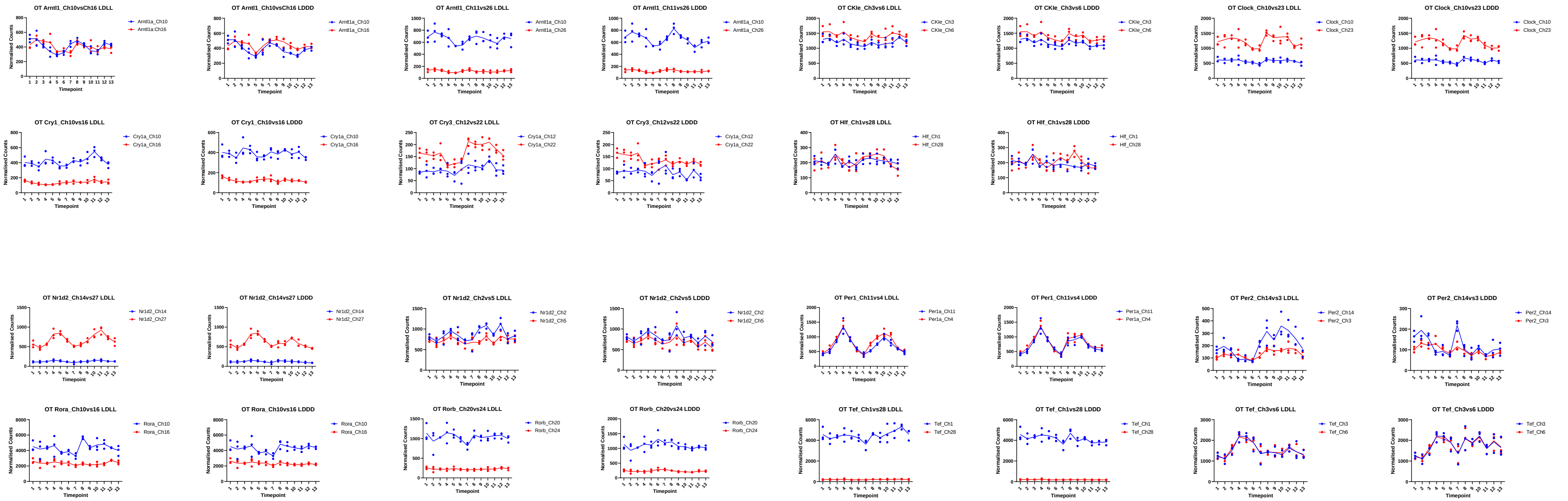

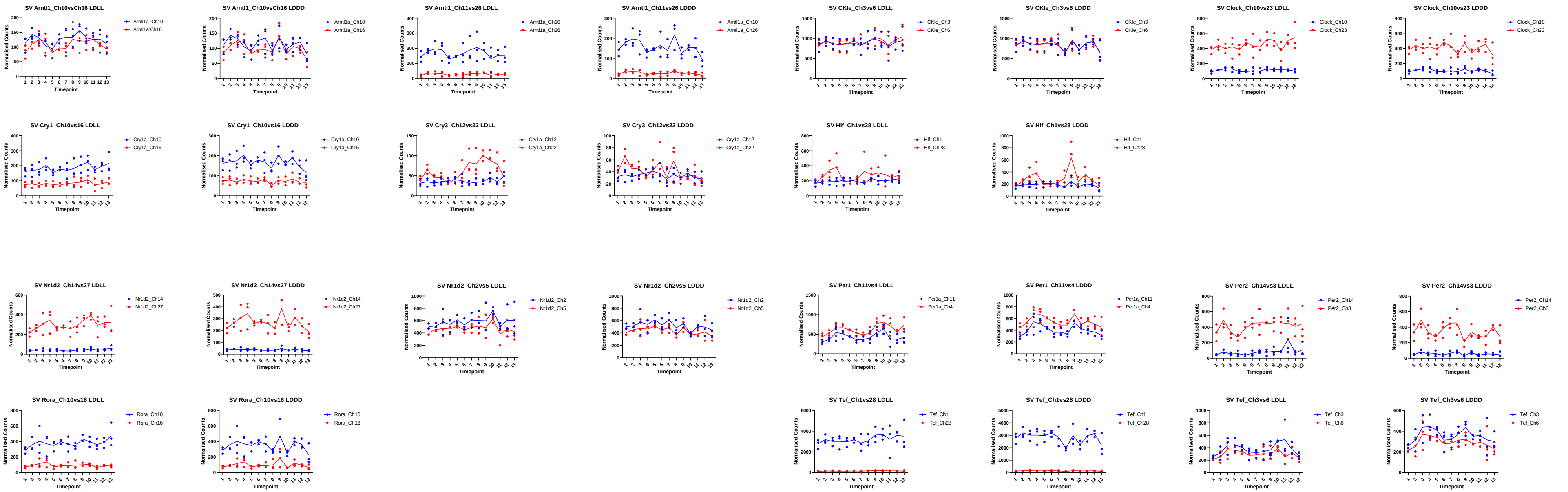

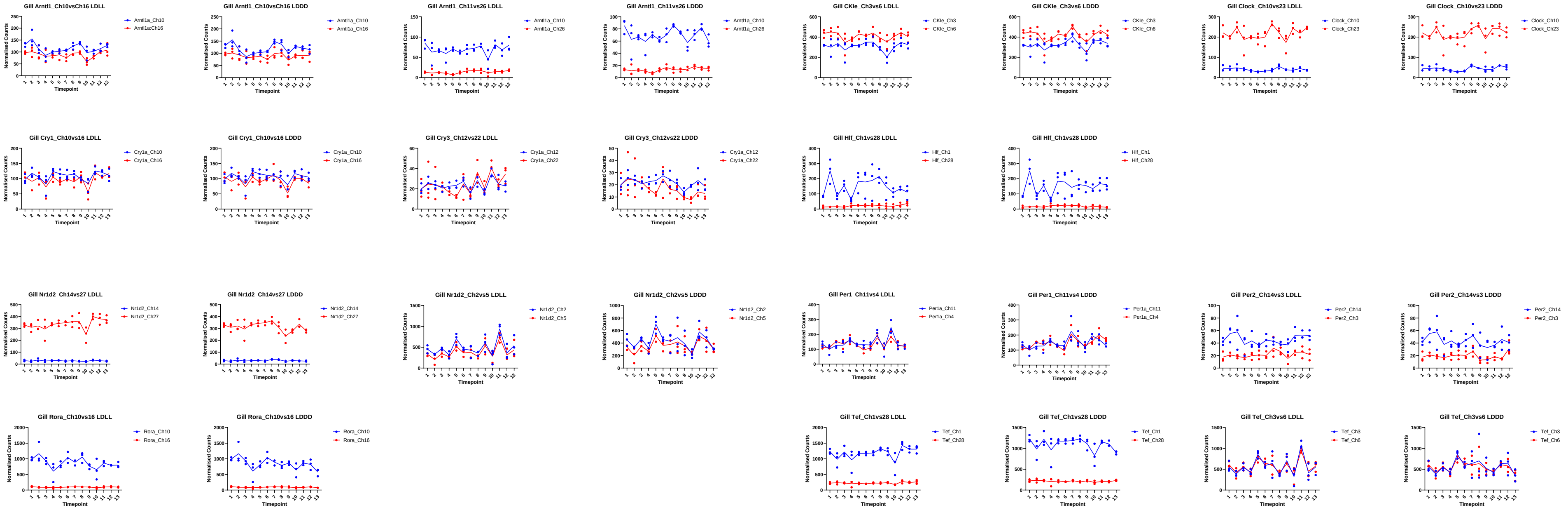
